# SiCEP3, a C-terminally encoded peptides from *Setaria italica*, promotes ABA import and signaling pathway

**DOI:** 10.1101/2021.01.30.428944

**Authors:** Lei Zhang, Yue Ren, Yiman Wan, Qian Xu, Guodong Yang, Shizhong Zhang, Jinguang Huang, Kang Yan, Chengchao Zheng, Changai Wu

**Author notes:** These authors contributed equally to this work (Lei Zhang, Yue Ren). **Corresponding authors** Chengchao Zheng, Changai Wu. **E-mails:** Lei Zhang, Yue Ren, Yiman Wan, Qian Xu, Shizhong Zhang, Guodong Yang, Jinguang Huang, Kang Yan, Chengchao Zheng, Changai Wu.

## Abstract

C-terminally encoded peptides (CEPs) are small peptides, typically post-translationally modified, and highly conserved in many species. CEPs are known to play roles in inhibition of plant growth and regulation of development, but the mechanisms are not well understood. In this study, we searched for CEP peptides in foxtail millet (*Setaria italica*). The 14 peptides we identified are divided into two subfamilies. The transcripts of most *SiCEP*s were more abundant in roots than in other tissues. *SiCEP3*, *SiCEP4,* and *SiCEP5* were also expressed at high levels in panicles. Moreover, expression of all *SiCEPs* was induced by biotic stress and phytohormones. *SiCEP3* overexpression and application of biosynthetic SiCEP3 both inhibited the growth of seedlings. In the presence of ABA, growth inhibition and ABA content of seedlings increased with the concentration of SiCEP3. Transcripts encoding two ABA transporters and one ABA receptor were induced by SiCEP3, ABA, and the two in combination. Further analysis revealed that SiCEP3 promoted ABA transport via NRT1.2 and ABCG40. In addition, SiCEP3, ABA, or the combination inhibited the kinase activities of CEP receptors CEPR1/2. Taken together, our results indicated that the CEP–CEPR module mediates ABA signaling by regulating ABCG40, NRT1.2, and PYL4 *in planta*.

**Highlight:** SiCEP3, a C-terminally encoded peptide, can promote ABA import and signaling pathway by inhibiting the kinase activity of its receptors under abiotic stress in *Setaria italica*.

## Introduction

Plant roots play vital roles in taking up water and nutrition to support plant growth, productivity, and survival in response to environmental stress (Hsu and Micallef, 2017). Root growth is dynamically regulated by a network of interacting hormone signals that coordinate cell division and differentiation (Lee et al., 2013). Auxin plays important roles in root development (Overvoorde et al., 2010), and other phytohormones regulate root growth through complex crosstalk with auxin signaling pathways (Koltai, 2015). High concentrations of abscisic acid (ABA) inhibit primary root growth (Ma et al., 2019). ABA signaling promotes ethylene production via phosphorylation of the C-terminus of 1-AMINOCYCLOPROPANE-1-CARBOXYLIC ACID SYNTHASE (ACS) and stabilization of ACSs by ABA-activated calcium-dependent protein kinase 4 (CPK4) and CPK11 (Luo et al., 2014). Ethylene signaling upregulates WEAK ETHYLENE INSENSITIVE2 (WEI2)/ANTHRANILATE SYNTHASE ⍰ 1 and WEI7/ANTHRANILATE SYNTHASE β1 to synthesize auxin and inhibit primary root growth (Stepanova and Alonso, 2005). ABA also promotes the production of reactive oxygen species (ROS) in mitochondria, reduces auxin accumulation, resulting in inhibition of primary root growth (He et al., 2012; Yang et al., 2014).

Recent studies revealed that small secreted CEPs (C-terminally encoded peptides) regulate root development in *Arabidopsis thaliana, Medicago truncatula*, *Nicotiana benthamiana*, *Samolous parviflorus*, and *Raphanus sativus* (Delay et al., 2013; Imin et al., 2013; Ohyama et al., 2008; Roberts et al., 2013; Yu et al., 2019b). In *Arabidopsis*, *AtCEP1* and *AtCEP3* decrease primary root growth and emerged lateral root number when these overexpressed or when the peptides they encode are applied to wild-type roots (Delay et al., 2013; Ohyama et al., 2008). In addition, the *Atcep3* mutant exhibits greater primary root growth and emerged lateral root number when grown under abiotic stress conditions (Delay et al., 2013). The identification of CEP receptors 1 and 2 (CEPR1 and CEPR2) in *Arabidopsis* suggested that sensing of root-derived CEP peptides by these shoot receptors mediates systemic nitrogen-demand signaling (Tabata et al., 2014). Application of CEP3 peptide decreases cell division, S-phase cell number, root meristematic cell number, and meristem zone (MZ) size in a dose- and CEPR1-dependent manner, especially under severe nutrient limitation (Delay et al., 2019). In *Medicago*, *MtCEP1* is expressed in primary root tips, vascular tissue, and lateral organ primordia, particularly under nitrogen (N) limitations (Imin et al., 2013). The *Medicago* COMPACT ROOT ARCHITECTURE2 (MtCRA2) receptor-like kinase is essential for inhibiting root growth in response to low N, leading to a repression of MtYUC2 auxin biosynthesis gene expression (Zhu et al., 2020). Together, these data suggest a relationship between CEP peptide signaling and auxin. Recently, we showed that CEPR2 participates in the ABA signaling pathway. Under normal conditions, CEPR2 phosphorylates ABA receptors (Yu et al., 2019b) and the ABA importer NPF4.6/NRT1.2 (Zhang et al., 2021) to promote their degradation and inhibit ABA signals. In the presence of ABA, CEPR2-medicated phosphorylation of NRT1.2 and ABA receptors is inhibited, maintaining their activities in ABA absorption and signaling. These observations suggest a relationship between CEP peptide signaling and ABA. However, it remains unclear whether ABA or abiotic stress are regulated by CEPs.

Foxtail millet (*Setaria italica*) is an important cereal crop that was domesticated from its wild ancestor, green foxtail millet (*Setaria viridis*) (Jia et al., 2013; Zhang et al., 2011). Both species are evolutionarily close to several bioenergy crops, including switchgrass (*Panicum virgatum*), napiergrass (*Pennisetum purpureum*), and pearl millet (*Pennisetum glaucum*), as well as major cereals such as sorghum and maize. Recently, the whole-genome sequences of foxtail millet and green foxtail millet have become available (Zhang et al., 2012). Both species have been proposed as C4 model plants (Lata et al., 2013). Along with progress in transformation techniques, the development of xiaomi, a mini foxtail millet with a life cycle similar to that of *Arabidopsis*, has increased the utility of foxtail millet as model C4 plant (Yang et al., 2020). Although studies of foxtail millet have recently advanced, the mechanisms of growth, organ development, and responses to biotic and abiotic stressors remain poorly understood.

In this study, to provide more information about the relationship between CEP peptides and ABA and abiotic stresses, we characterized CEP peptides in foxtail millet. We found that all *SiCEPs* were highly expressed in roots, and that a cluster of genes was also highly expressed in panicles. All *SiCEPs* were induced by salt stress, osmotic stress, or treatment with ABA or IAA. Both application of SiCEP3 application and overexpression of *SiCEPs* in hairy-roots repressed root and shoot growth in foxtail millet seedlings. Furthermore, SiCEP3 promoted the transport of ABA by activating the expression of genes encoding ABA transporters, such as ABCG40 and NPF4.6/NRT1.2. These findings provide novel insights into the role of CEP peptide signaling pathways in regulating plant growth and abiotic stress responses.

## Materials and Methods

### Plant growth and treatment

The plant materials used in this study were foxtail millet (*Setaria italica*) cultivar Yugu1, *Arabidopsis* Col-0, and related mutants, which have been used in previous studies of our lab, in the Col-0 background. Yugu1 seeds were sterilized with sodium hypochlorite, washed three times, and germinated for 3 d. The 3-day-old seedlings were cultured for 10 days or 2 months with Hoagland solution in greenhouse conditions under a 16/8 h light/dark period at 26 ± 1°C (Le Pioufle et al., 2019). For stress treatment, 10-day-old seedlings of Yugu1 were treated with Hoagland solution containing 200 mM NaCl, 100 μM ABA, 100 μM IAA, and 20% polyethylene glycol 6000 (PEG6000) for 0, 6, 12, or 24 h (Li et al., 2016). The roots and shoots were collected separately, rapidly frozen in liquid nitrogen, and subjected to RNA extraction. For tissue expression analysis, the roots, stems, leaves and panicles from 2-month-old seedlings of Yugu1 were collected separately and rapidly frozen in liquid nitrogen. For exogenous application of CEP and ABA, either alone or in combination, the uniformly developed 3-day-old seedlings of Yugu1 were transferred to Hoagland solution with or without 0.5, 1, or 2 μM CEP, 7.5 μM ABA, or the combination, and cultured for an additional 7 days (Liu et al., 2020). For the seedlings with transgenic hairy roots, the 3-day-old shoots were incubated with *Agrobacterium* K599 containing Ubi:SiCEP3 or empty vector and cultured on MS with Timentin (Sangon, Shanghai, China). Culture methods for *Arabidopsis* materials were described previously (Chen et al., 2014). Phenotypes were photographed, and root or shoot lengths were measured. Toot samples were used to measure ABA and detect expression of related genes.

### Identification of CEP genes in Foxtail millet

To find *SiCEP* genes, the CEPs of *Arabidopsis* were used as query sequences to BLAST the foxtail millet genome in Phytozome V12 (https://phytozome.jgi.doe.gov) (Roberts et al., 2013). Signal peptides were predicted using the signalP5.0 software (http://www.cbs.dtu.dk/services/SignalP/). The structures of CEP genes were visualized using GSDS (Gene Structure Display Server: http://gsds.cbi.pku.edu.cn/). Isoelectric point and molecular weight were predicted using the ExPASy Proteomics Server (http://expasy.org/). Evolutionary trees and mature peptides were generated using MEGA7.0. The cis-elements on the promoters of CEP genes were predicted using PlantCARE (http://bioinformatics.psb.ugent.be/webtools/plantcare/html/). Three-dimensional models of CEP proteins were generated by I-TASSER (https://zhanglab.ccmb.med.umich.edu/I-TASSER/) (Roy et al., 2010)

### RNA Extraction and RT-qPCR Analysis

Total RNA was extracted using the Plant RNA Purification Reagent (Invitrogen, Carlsbad, CA, USA) as described in previous studies (Yu et al., 2019b). First-strand cDNA was synthesized from 2 μg of total RNA with the *Evo M-MLV* RT Kit with gDNA Clean for qPCR II AG11711 (Accurate Biotechnology (Hunan)Co., Ltd). RT-qPCR was performed using ChamQ Universal SYBR qPCR Master Mix (Vazyme Biotech, Nanjing, China) with a three-step program on a CFX96TM Real-Time PCR Detection System (Bio-Rad, Hercules, CA, USA) (Xu et al., 2018). The 18S ribosomal RNA and *SiACT7* (Li et al., 2016) were used as controls. Three biological replicates were performed. Primer sequences are provided in Supplementary Table S1.

### SiCEP3 uptake assays

SiCEP3 (GWMPDGSVPSPGVGH) (75% purity) was chemically synthesized by Sangon (Shanghai, China, http://www.sangon.com/) and N-terminally labeled with fluoresce in isothiocyanate (5-FITC) by Link Biotech. Ten-day-old Yugu1 seedlings were incubated with 5-FITC-SiCEP3 (10 μM) in MS at 25°C for the indicated times. The seedlings were washed three times for 5 min by gentle shaking in ddH_2_O, and then photographed on a LUYOR-3109D with a excitation at 495 nm and emission at 545 nm.

### Generation of SiCEP3-overexpressing roots

Full-length CDS of SiCEP3 was amplified and cloned into pTCK303 vector under the control of the ubiquitin promoter (Du et al., 2016). The recombinant plasmid was transformed into *A. tumefaciens* K599, which was cultured to an OD_600_ = 1.0 with continuous shaking. The culture was then centrifuged at 6,000 g for 5 min, and MS liquid medium was added to resuspend the bacteria at a final concentration of OD_600_ = 1.0. Three-day-old shoots were incubated with *A. tumefaciens* containing ubi:SiCEP3 at 28°C for 10–20 min with continuous shaking, and then transferred to MS solid medium supplemented with carbenicillin (250 mg L^−1^) for one night to induce root formation (Bahramnejad et al., 2019). Three days later, transgenic seedlings were transferred to MS liquid medium with or without 7.5 μM ABA for 7 d. The phenotype was photographed and the root length was measured. The root samples were used to measure ABA content and expression of related genes.

### *In vitro* kinase assays

The CDS sequences encoding the kinase domain of CEPR1 (AT5G49660, amino acids 614–966) and CEPR2 (AT5G49660, amino acids 642–977) were amplified and cloned into pET30a. Transformation, induction, and purification of fusion proteins were performed as described previously (Yu et al., 2019b). *In vitro* kinase assays were carried out using purified CEPR1^KD^-His or CEPR2^KD^-His. We combined 50 μg of CEPR1^KD^-His or CEPR2^KD^-His in 50 μL of reaction buffer (25 mM HEPES, pH 7.2, 1 mM DTT, 50 mM NaCl, 2 mM EGTA, 5 mM MgSO4, and 50 μM ATP). Reaction mixture with or without ABA/SiCEP3 was incubated at 22°C for 0–2 h and terminated by addition of 5× loading buffer (Zhang et al., 2021). Sample without adding ATP reaction was used as the negative control. Proteins were then fractionated on SDS-PAGE and Mn^2+^-Phos-tag-PAGE (50 μM Phos-tag and 100 μM Mn^2+^). After incubation for 2 h, 0.01 U/μL calf intestinal alkaline phosphatase (CIAP; Promega, Madison, WI, USA) was added to the reaction buffer, and the sample was incubated for 30 min at 37 °C to remove the phosphoryl group(s) from CEPR1^KD^-His or CEPR2^KD^-His. Phosphorylation level was detected by western blotting using mouse anti-His monoclonal antibody (HT501-01, TransGen Biotech).

## Results

### Identification and analysis of CEP genes in *Setaria italica*

To identify the canonical CEP genes, we scanned the foxtail millet genome assembly (*Setaria italica* v2.2 in JGI Phytozome, https://phytozome.jgi.doe.gov) was scanned for open reading frames (ORF) with CEP domains similar to 15 CEP domain sequences from *Arabidopsis*. Query sequences with E-value < 1 were selected, resulting in identification of 14 *CEP* ORFs were identified. The chromosomal distribution of *SiCEP* genes in foxtail millet was uneven: six of nine chromosomes contained *SiCEP* genes (Fig. 1A). Among them, chromosome 3 and chromosome 9 each had four *SiCEP* genes; chromosomes 2 and 5 each had two *SiCEP* genes, and chromosomes 4 and 6 each only have one gene each. Based on the chromosomal localization (i.e., starting with chromosome 2 and proceeding through chromosome 9), we named these genes as *SiCEP1* to *SiCEP14* (Fig. 1A). Structural analysis revealed that all *SiCEP* genes have only one exon, with no introns (Fig. S1A). Except for *SiCEP1* and *SiCEP14*, which have no untranslated regions (UTRs), all other *SiCEP* genes have UTRs of different lengths (Fig. S1A). Three *SiCEP* gene clusters were found in chromosome 2, 3 and 9, containing *SiCEP1–2*, *SiCEP*3–5, and *SiCEP11–14* (Fig. 1A). Signal peptide prediction revealed that all SiCEPs have an N-terminal signal peptide with about 20 amino acids (Fig. S1B, Table S2). Analysis with MEGA7.0 revealed that *SiCEP*s were similar to *AtCEP*s, which c divided into two distinct groups: Group I, with 8 members, and Group II with 6 members (Fig. 1B). Interestingly, all SiCEP peptide precursors had only one CEP domain (Fig. S1B), whereas many genes across species contain multiple CEP domains (Ogilvie et al., 2014; Yu et al., 2019b). The CEP domain formed a conserved helical structure at the C-terminus of each predicted protein model (Fig. S1C). Analysis of physical and biochemical information (https://web.expasy.org) analysis revealed that SiCEP precursor proteins contained between 76 and 111 amino acids and had isoelectric points (pI) ranging from 4 to 12; the mature peptides each contained about 15 amino acids (Table S2). The conserved 15–amino acid CEP domains were completely identical between SiCEP1 and SiCEP2, and between SiCEP12 and SiCEP13, but varied among members of SiCEP3-5 and SiCEP11-14 (Table S2). Due to the low concentration in planta, peptide hormones often appear in gene clusters due to gene duplication, allowing amplification or neo-functionalization (Matsubayashi, 2014). Therefore, the *SiCEP*1–2, *SiCEP*3–5, and *SiCEP*11–14 genes might have both redundant and new functions.

**Figure 1.**
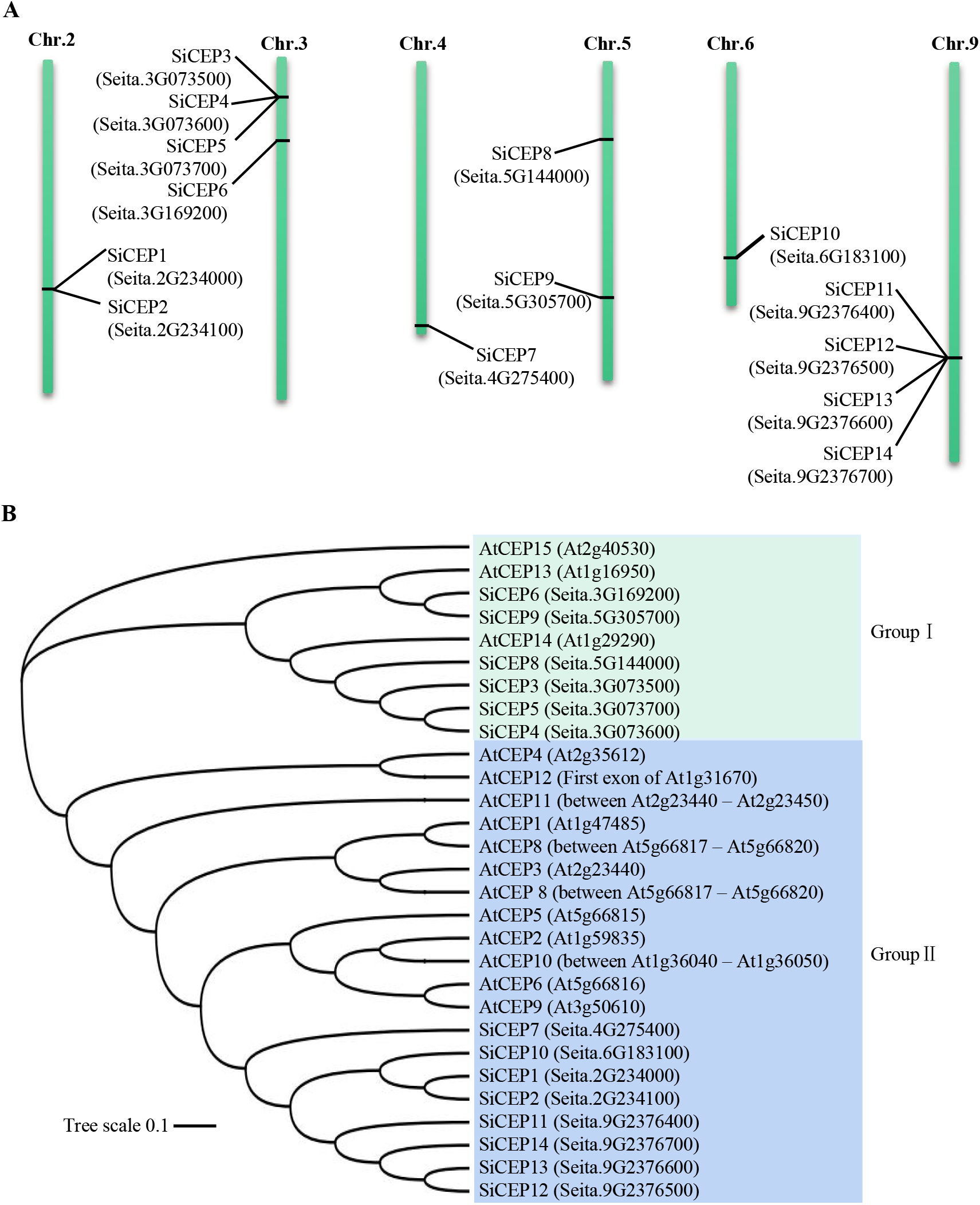
Identification and analysis of CEP precursor genes in *Setaria italica*. (A) Distribution of *SiCEP* genes on chromosomes of *Setaria italica*. (B) Phylogenetic analysis of CEPs of *Arabidopsis* and *Setaria italica*. The phylogenetic tree, based on CEP domains, was created using MEGA7 by the neighbor-joining method. Bootstrap tests are indicated on the tree.

### Expression patterns of SiCEP genes in different tissues

To explore the spatial expression of *SiCEP* genes in roots, stems, functional leaves, and panicles from 2-month-old Yugu1 seedlings, we performed quantitative RT-PCR (RT-qPCR). The results revealed that the *SiCEP* genes exhibited relatively conserved expression patterns in terms of tissue specificity and levels (Fig. 2A). All *SiCEP* genes were highly expressed in roots ((Fig. 2A, Fig. S2A), similar to *CEP*s in *Arabidopsis thaliana* (Ohyama et al., 2008), *Medicago sativa* (Imin et al., 2013), and *Malus hupehesis* (Yu et al., 2019a). These observations suggested that *SiCEP*s regulate root development or control substance absorption in roots. Surprisingly, almost all *SiCEP* genes except for *SiCEP8* and *SiCEP9* were expressed at very low levels in stems and leaves. *SiCEP9* also showed high expression in stems and leaves, as well as in panicles, indicating its role in regulating multiple organ development. Interestingly, one gene cluster composed of SiCEP3, SiCEP4 and SiCEP5 was also expressed at high levels in panicles, indicating possible roles of these three genes in the development of panicles, flowers, seeds, or other related organs. The tissue-specific and overlapping expression patterns of *SiCEP*s in various tissues indicated that these genes (and their clusters) may play similar or different roles.

**Figure 2.**
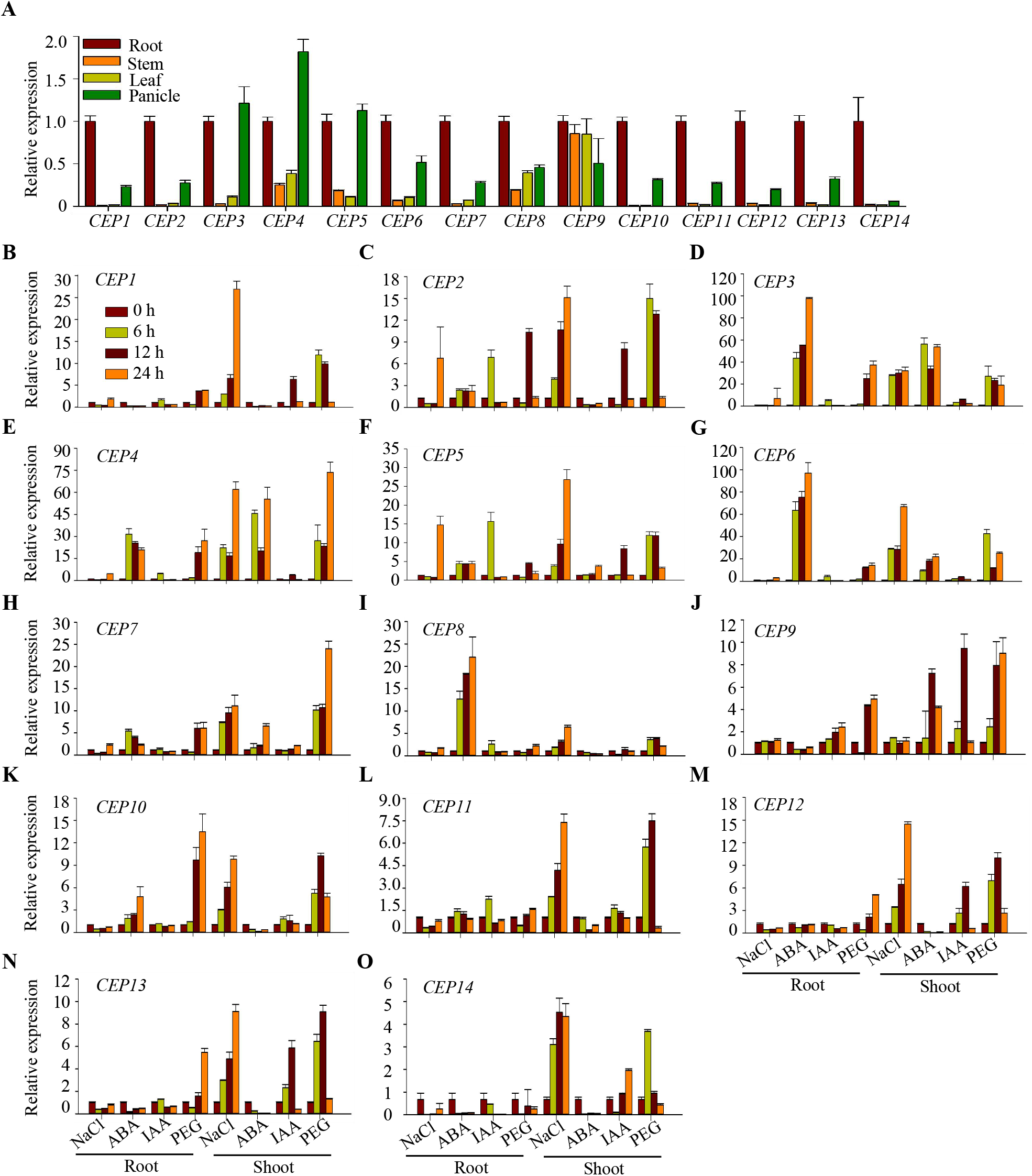
Expression patterns of *SiCEP*s in Yugu1 seedlings under abiotic stresses. (A) Expression levels of *SiCEP1*–*SiCEP14*, detected by qRT-PCR, in root, stem, leaf, and panicle of 2-month-old Yugu1 seedlings. *ACT7* and 18s rRNA were used as internal controls. Error bars indicate SEM (n = 3), Experiments were performed in biological triplicate. (B)–(O) Expression levels of *SiCEP1*–*SiCEP14*, detected by qRT-PCR, in roots and shoots of 10-day-old Yugu1 seedlings after treatment with 200 mM NaCl, 100 μM ABA, 100 μM IAA, and 20% PEG6000 for 0, 6, 12, or 24 hours. Error bars indicate SEM (n = 3), p < 0.05. One-way ANOVA Duncan’s test was used for statistical analysis. Experiments were performed in biological triplicate. Values are shown as fold changes relative to mock-treated samples and normalized against 18s rRNA.

### Transcriptional profiles of *SiCEP* genes under abiotic stresses

To elucidate the potential involvement of *SiCEP* genes in abiotic stresses, we analyzed the cis-elements in the promoter 1500 bp upstream of the initiation codon of each *SiCEP* gene (Fig. S2B). The results revealed multiple hormone-responsive elements, including the jasmonic acid response elements TGACG, CGTCA, and the TATC-box for; the auxin response elements TGA-element and AuxRR-core, the ABRE (ABA-responsive element), the ERE (ethylene response element), and the salicylic acid response element TCA (Fig. S2B and C). Together, these results suggested that *SiCEP* genes respond to multiple phytohormones.

Furthermore, we detected transcript levels of *SiCEP* genes by qRT-PCR in 7-day-old Yugu1 seedlings treated with phytohormones ABA and IAA, as well as NaCl and PEG6000 (Fig. 2B–2O). Among 14 *SiCEP* genes, the upregulation of *SiCEP3*, *SiCEP4*, *SiCEP5*, and *SiCEP6* (up to 100-fold) was higher than that of other *SiCEP* genes (Fig. 2D–G), whereas the upregulation of SiCEP11, SiCEP13, and SiCEP14 was lower than 10-fold under stress conditions (Fig. 2 L, N, and O). Under salt stress, expression of all *SiCEP* genes was significantly induced in shoots (Fig. 2B - 2O). Similar induction patterns were detected in response to PEG6000. When treated with ABA, expression of *SiCEP3*, *SiCEP4*, and *SiCEP6* was significantly upregulated in both roots and shoots, whereas *SiCEP8* gene in roots and others were induced less than 10-fold, or were inhibited, in shoots and roots. By contrast, expression of *SiCEP* genes was not significantly induced by IAA. Interestingly, *SiCEP* genes clustered on chromosomes exhibited similar inducible expression patterns of inducible expression. For example, SiCEP11, SiCEP12, and SiCEP14 were induced by salt and osmotic stresses (Fig. 2L, M, and O), whereas SiCEP3 and SiCEP4 were significantly upregulated by ABA, salt and osmotic stresses (Fig. 2D and F). These results indicated that *SiCEP*s might play important roles in response to ABA-mediated abiotic stresses, including drought and excess salt.

### SiCEP3 promotes ABA transport and signaling

To investigate the relationship between CEPs and ABA, we treated Yugu1 seedlings with synthesized mature SiCEP3 peptide (GWMPDGSVPSPGVGH) (Table S2) in the presence or absence of ABA. The growth of Yugu1 seedlings was inhibited by external SiCEP3, as reflected by significant shortening of roots (Fig. 3A and B). External ABA treatment resulted in similar growth inhibition (Fig. 3A and B). In the presence of ABA, seedling growth worse and shorter roots were abtained with the concentrations of SiCEP3 increasing (Fig. 3A and B). Fluorescence increased with the duration of SiCEP3 application (Fig. 3C), indicating that more and more SiCEP3 was taken up by Yugu1 seedlings. The increase in ABA content resulting from elevated concentrations of SiCEP3 in the presence of ABA (Fig. 3D) indicated that SiCEP3 promotes ABA uptake. When SiCEP3, ABA, or the combination was applied for a long period of time, plant growth was inhibited, and roots were shortened (Fig. S3A and S2B). Moreover, expression of *SiABCG40 (Seita.5G221600)*, *SiNRT1.2 (Seita.4G231500)*, and *SiPYL4 (Seita.3G207900)*, which encode the ABA transporter and ABA receptor, respectively, was induced by SiCEP3, ABA, or the combination (Fig. 3E, 3F and 3G). These data indicated that SiCEP3 promotes ABA uptake and ABA signaling.

**Figure 3.**
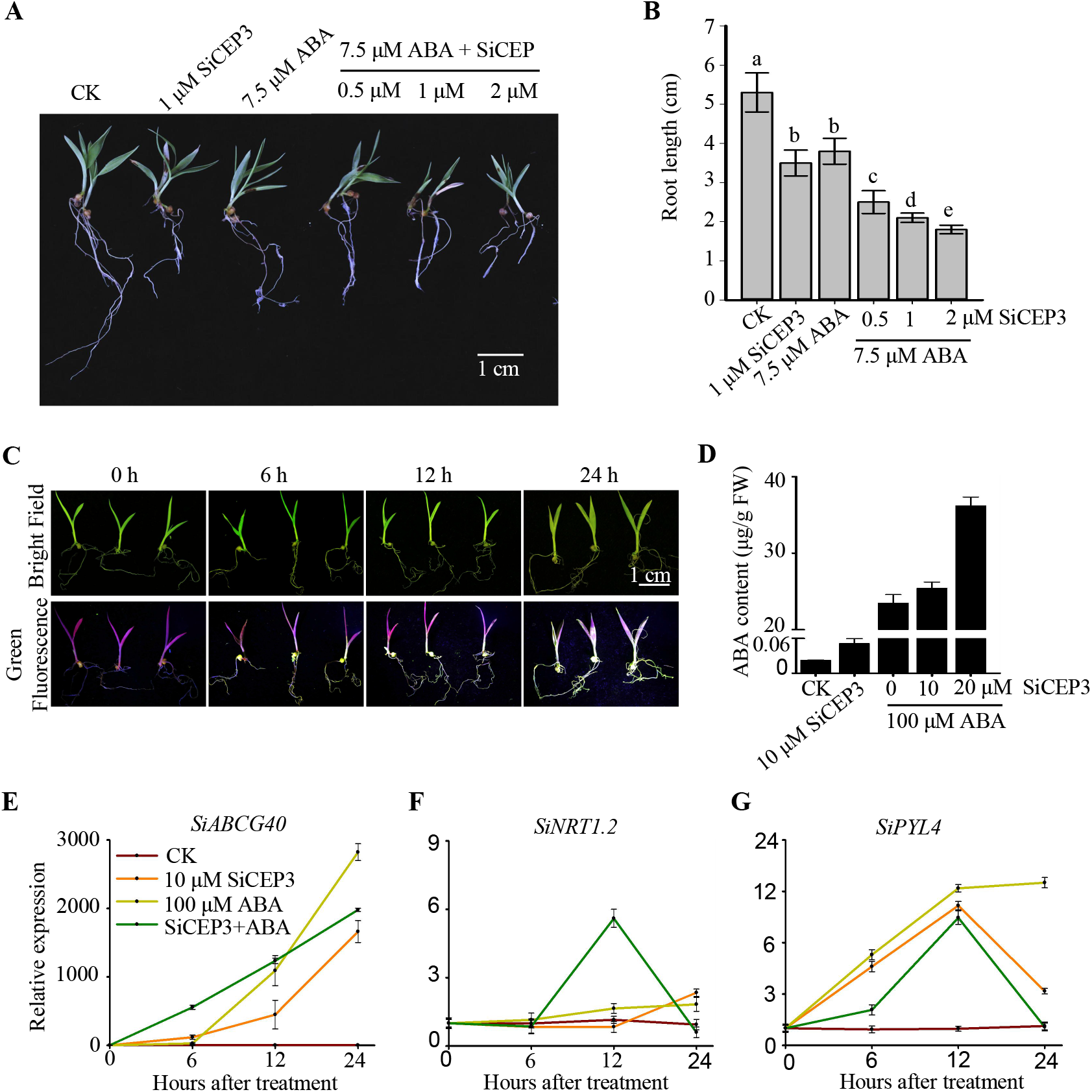
External application of SiCEP3 affects plant growth, ABA content, and gene expression in Yugu1 seedlings. (A) Effects of SiCEP3 on Yugu1 seedlings treated with or without ABA. Three-day-old Yugu1 seedlings were transferred onto MS medium containing 1 μM SiCEP3 and 7.5 μΜ ABA or the combine for additional 7 d. (B) Primary root length and plant height of the seedlings in (A). Error bars indicate SEM (n = 3). Bars labeled with different lowercase letters are significantly different from one another (P < 0.05; one way ANOVA). (C) SiCEP3 uptake by Yugu1 seedlings. Uptake was monitored by fluorescence emission of 5-FITC-CEP3. The experiment was repeated three times independently. (D) Effects of SiCEP3, ABA, and the combination on ABA content. Ten-day-old Yugu1 seedlings were grown on MS containing 10 μM SiCEP3, 100 μM ABA, 10 μM SiCEP3+100 μM ABA, or 20 μM SiCEP3+100 μM ABA for 6 h. Error bars indicate SEM (n = 3). Bars labeled with different letters in lowercase are significantly different from one another (P < 0.05; one way ANOVA). (E)–(G)Transcript levels of genes encoding ABA transporters *SiABCG40* and *SiNRT1.2*, and ABA receptor *SiPYL4* in Yugu1 seedlings under normal conditions or with 10 μM SiCEP3, 100 μM ABA, or the combination for 0, 6, 12, and or 24 h. Transcript levels were measured by qRT-PCR

To further illustrate the effects of SiCEP3 on the ABA uptake and signaling, we established the *Agrobacterium tumefaciens*–mediated hairy-root transformation technique with high transformation efficiency (up to 76%). Expression of SiCEP3 in transgenic seedlings was about 6-fold higher than in empty-vector transgenic seedlings (Fig. 4B). SiCEP3 transgenic seedlings had shorter roots than empty-vector transgenic seedlings in the presence and absence of ABA (Fig. 4A and C). SiCEP3 transgenic seedlings had higher ABA levels than empty vector transgenic seedlings under both conditions (Fig. 4D). These data suggested that SiCEP3 slightly promotes ABA biosynthesis and significantly promotes ABA absorption. This idea was further confirmed by the upregulation of *SiABCG40*, *SiNRT1.2* and *SiPYL4* and anthocyanin accumulation in SiCEP3 transgenic seedlings relative to empty-vector transgenic seedlings (Fig. 4A, E, F and G).

**Figure 4.**
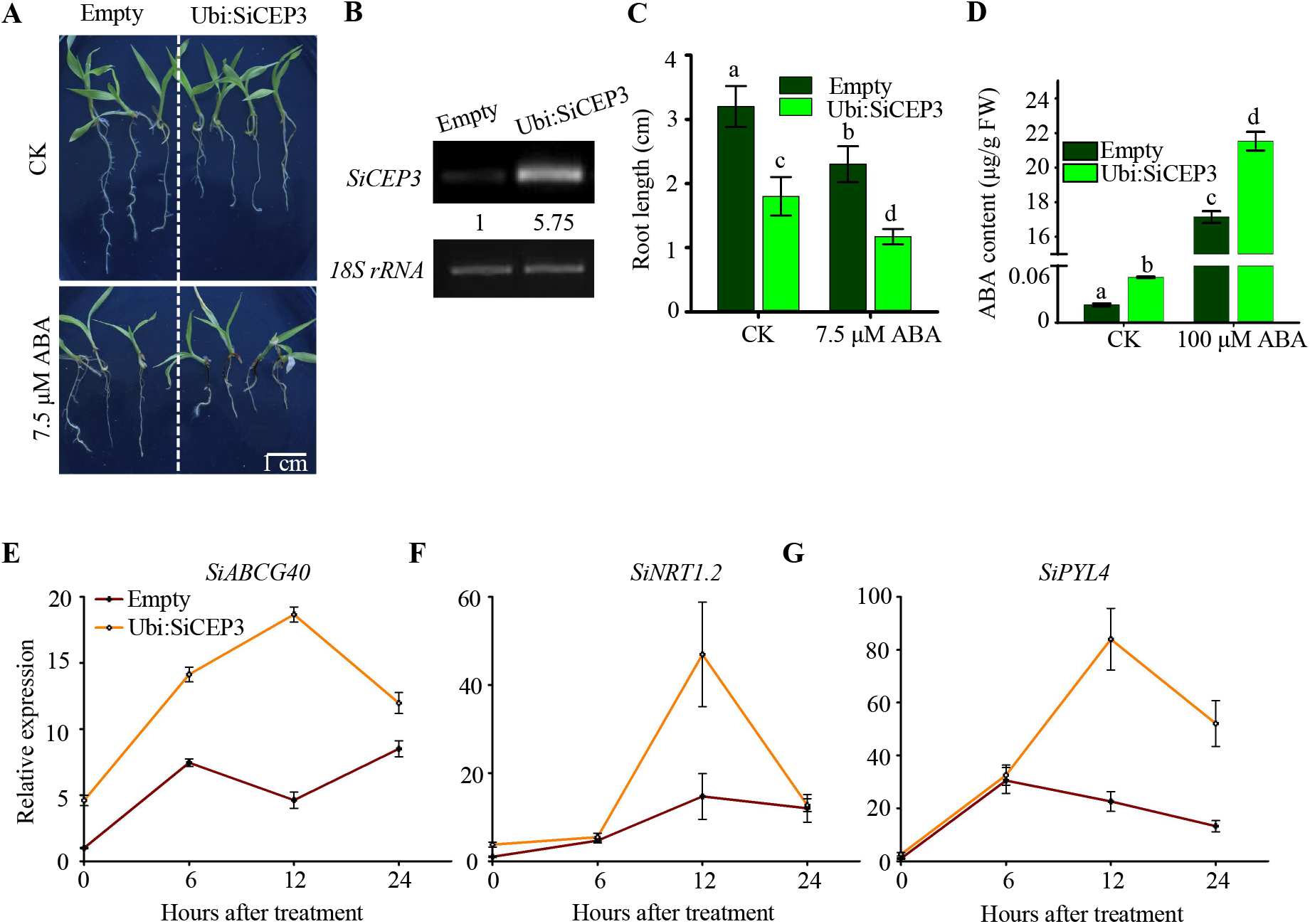
Overexpression of SiCEP3 in hairy roots inhibits plant growth, ABA content, and gene expression in Yugu1 seedlings. (A) Images of transgenic Yugu1 seedlings treated with or without 7.5 μM ABA for 7 d. The data represent the average of three biological replicates. (B) Expression levels of *SiCEP3,* as determined by RT-PCR, in transgenic and control hairy roots. Levels of *SiCEP3* mRNA were normalized against 18S rRNA. (C) Primary root length of seedlings in (A). Error bars indicate SEM (N = 3). Bars labeled with different lowercase letters are significantly different from one another (P < 0.05; one way ANOVA). (D) ABA contents of 14-day-old empty-vector Ubi:SiCEP3 transgenic seedlings grown on MS with or without 10 μM SiCEP3, 100 μM ABA, or the combination for 6 h. Error bars indicate SEM (n = 3). Bars labeled with different letters in lowercase are significantly different from one another (P < 0.05; one way ANOVA). (E)–(G) Transcript levels of *SiABCG40*, *SiNRT1.2*, and *SiPYL4* in *SiCEP3* transgenic hairy roots, as determined by qRT-PCR. Hairy roots transformed with the empty vector were used as controls.

Next, we generated the *nrt1.2abcg40* double mutant by crossing *nrt1.2* with *abcg40*. When SiCEP3, ABA, or the combination was applied to *Arabidopsis* seedlings of wild-type (WT) and ABA transporter mutants, root growth of wild-type *Arabidopsis* seedlings was inhibited by external SiCEP3 and ABA, and was aggregated by the combination of SiCEP3 and ABA (Fig. 5A and B), indicating a conserved function of SiCEP3 in *Arabidopsis*. The *nrt1.2abcg40* and *pyr1/pyl24* seedlings exhibited no growth difference relative to WT seedlings after SiCEP3 treatment, whereas both mutants had longer roots than wild-type under ABA or combination treatment (Fig. 5A and B). A similar response was observed in seedlings harboring loss-of-function mutations in genes involved in ABA signaling, including *abi3* and *abi5* (Fig. S4A and B). However, loss-of-function mutations in genes involved in ABA biosynthesis, including *aba1*, *aba2* and *nced3*, were insensitive to SiCEP3 but exhibited different their response to ABA or combination treatment (Fig. S4A and B). The ABA content of WT seedlings increased in the presence of ABA and SiCEP3, whereas that of *nrt1.2abcg40* seedlings was lower than in WT (Fig. 5C). Expression of *AtABCG40*, *AtNRT1.2*, and *AtPYL4* was induced by SiCEP3, ABA, or the combination (Fig. 5D, E and F). These observations indicated that SiCEP3 mainly promotes ABA uptake, at least through NRT1.2 and ABCG40, which in turn leads to accumulation of ABA in certain tissues to activate ABA signaling.

**Figure 5.**
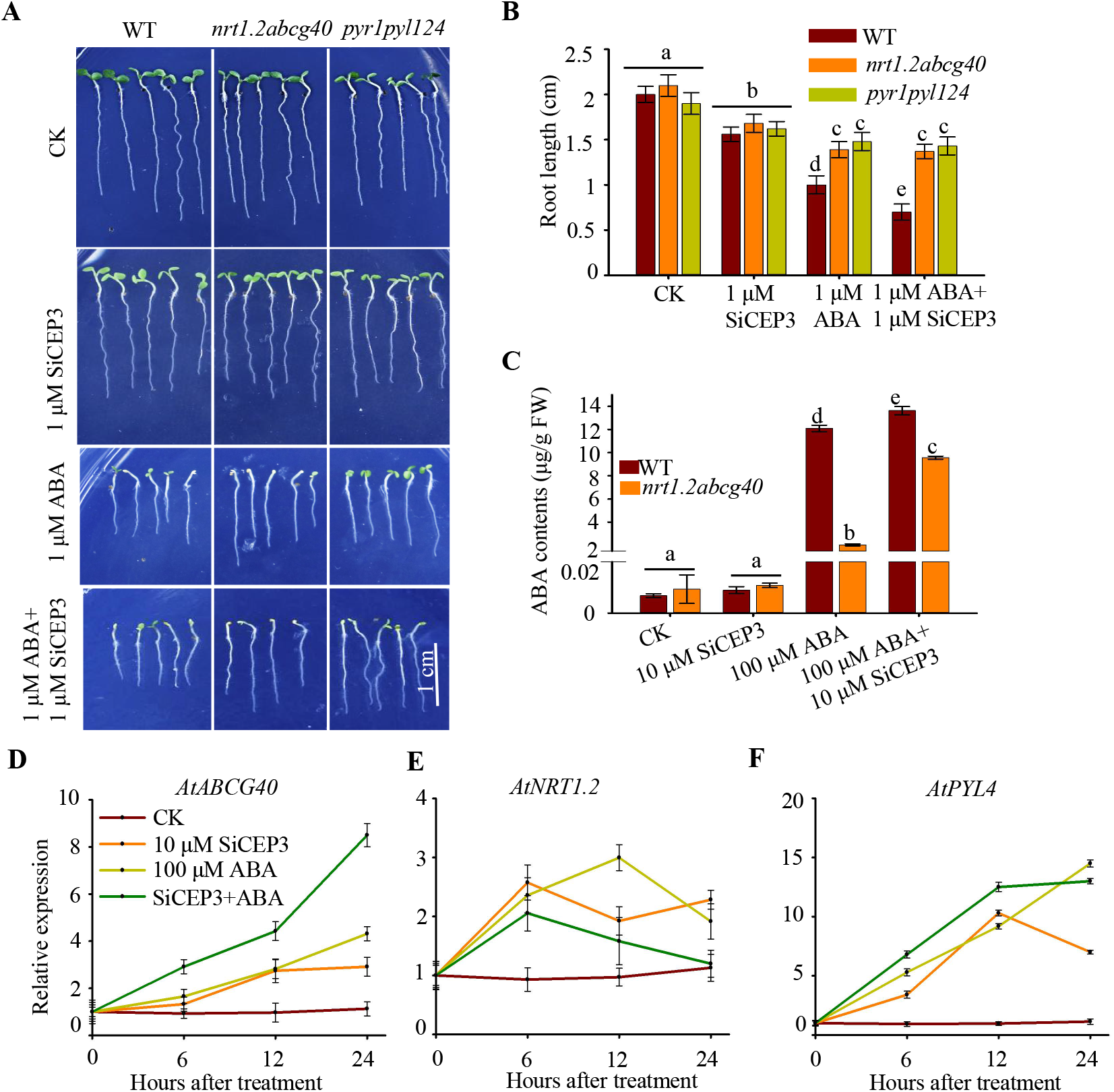
SiCEP3 inhibits ABA transport and signaling in *Arabidopsis*. (A) Phenotypes of WT, *nrt1.2abcg40*, and *pyr1/pyl124* mutant seedlings grown on 1/2 MS with or without 1 μM SiCEP3, 1 μM ABA, or the combination for 7 d. (B) Primary root length of seedlings shown in (A). Error bars indicate SEM (n = 3). Bars labeled with different lowercase letters are significantly different from one another (P < 0.05; one way ANOVA). (C) ABA contents of 7-day-old WT and *nrt1.2abcg40* mutant seedlings grown on 1/2 MS with or without 10 μM SiCEP3, 100 μM ABA, or the combination for 6 h. Error bars indicate SEM (n = 3). Bars labeled with different letters in lowercase are significantly different from one another (P < 0.05; one way ANOVA). (D)–(F) Transcript levels of *AtABCG40*, *AtNRT1.2,* and *AtPYL4*, determined qRT-PCR in *Arabidopsis* seedlings grown on 1/2 MS with or without 10 μM SiCEP3, 100 μM ABA, or the combination for 0, 6, 12, or 24 h.

### Both SiCEP3 and ABA inhibit the kinase activity of CEPR1 and CEPR2

CEPR1 and CEPR2 are CEP receptors (Tabata et al., 2014). Previously, we showed that the stability and activity of ABA receptor PYL4 and ABA importer NRT1.2 were regulated by CEPR2-mediated phosphorylation modification, and that interaction between CEPR2 and NRT1.2 or PYL4 does not depend on ABA (Yu et al., 2019b; Zhang et al., 2021). We hypothesized that the kinase activities of CEPR1 and CEPR2 are regulated by CEPs or ABA. To test this possibility, we performed *in vitro* kinase assays. After 2 h, we observed autophosphorylation or the kinase domains of CEPR1 and CEPR2 (Fig. 6A). When CIAP, CEP3, ABA, or the combination was added, phosphorylation of the kinase domain of CEPR1 and CEPR2 decreased or disappeared (Fig. 6A). These results indicated that CEP3 and ABA inhibit the kinase activities of CEPR1 and CEPR2.

**Figure 6.**
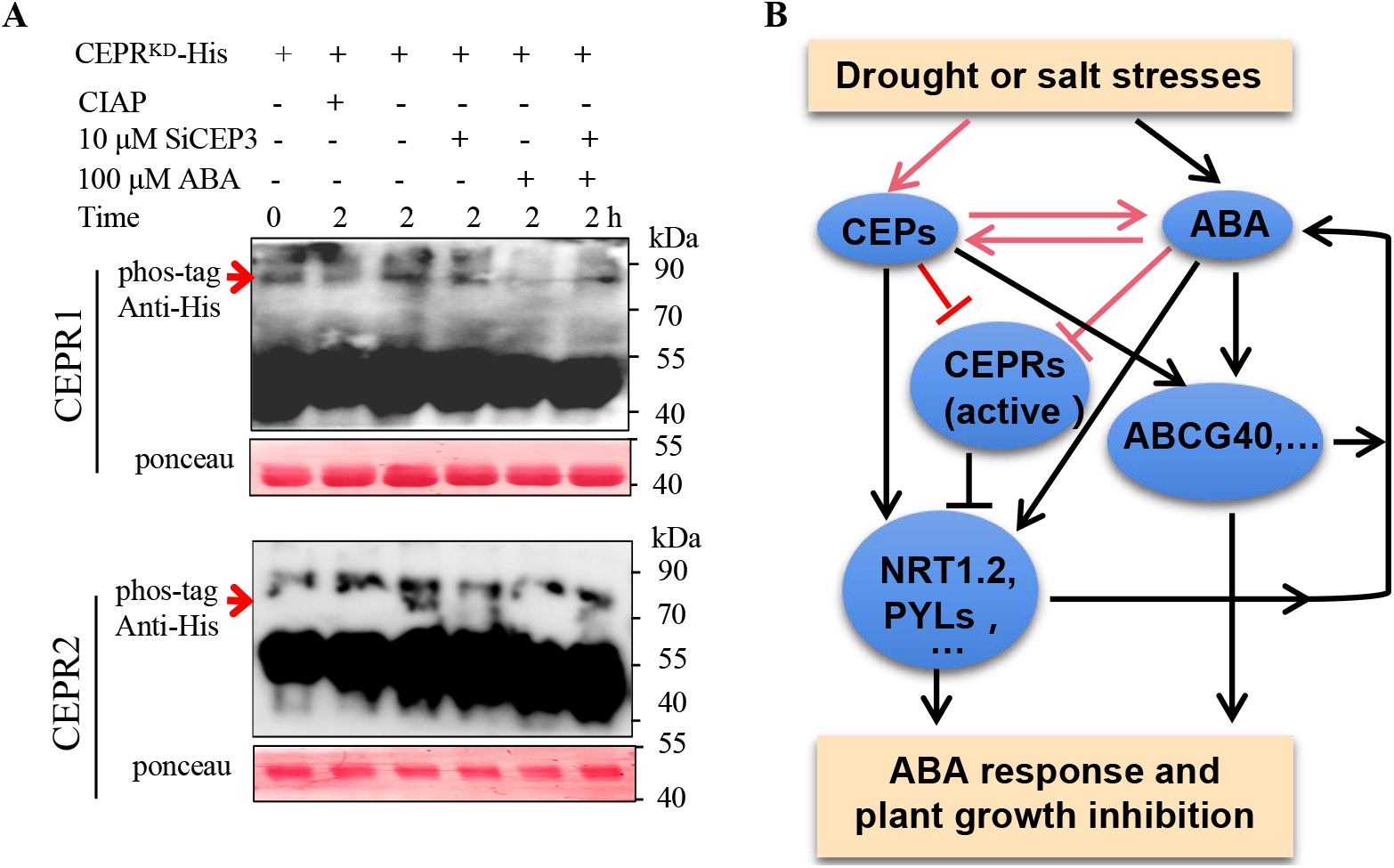
SiCEP3 inhibits the kinase activities of AtCEPR1 and AtCEPR2 and promotes ABA absorption and signaling. (A) *In vitro* kinase assays were performed using the kinase domains of AtCEPR1 and AtCEPR2 in the presence of CIAP, SiCEP, ABA, or the combination for 0 or 2 h. Autophosphorylation status and protein levels of AtCEPRs were detected by Phos-tag (upper) and western blotting (lower), respectively. Red arrows indicate the autophosphorylated kinase domain of AtCEPR1 or AtCEPR2. (B) CEP-mediated abiotic stress response in plants. Under normal conditions, low levels of CEPs and ABA are present within cells. CEPRs have high phosphorylation activity and phosphorylate NRT1.2, PYLs, and other related proteins to promote their degradation, thereby inhibiting transduction of ABA signal. Consequently, plants grow and develop normally. Under abiotic stresses, CEPs and ABA are highly induced and bound by their receptors. CEP-bound CEPRs lose their kinase activity, and hence cannot phosphorylate NRT1.2, PYLs, and other related proteins. Consequently, these proteins remain highly active, transporting ABA or transducing the ABA signal, leading to inhibition of plant growth.

Taken together, these findings provide novel information about the CEP–CEPR-mediated ABA response in plants. Salt and osmotic stresses that induce ABA accumulation, as well as ABA, induce expression of CEPs and ABA biosynthesis in foxtail millet seedlings. The CEPs are transported out of cells and bound by their receptors, CEPR1 and CEPR2. This, in turn, inactivates the kinases of CEPR1 and CEPR2 and maintains the stability and activity of ABA transporters and receptors to active ABA transport and signaling in response abiotic stresses, ultimately leading to plant growth inhibition (Fig. 6B). However, CEPS can also inhibit plant growth in an ABA-independent manner.

## Discussion

Small secreted CEPs have pleiotropic effects on plant growth in *Arabidopsis thaliana, Medicago* truncatula, and other plant species, influencing root growth, lateral root development, panicle and seed development, and responses to abiotic stresses, including low-nitrogen, salt, and mannitol stresses (Delay et al., 2013; Imin et al., 2013; Ohyama et al., 2008; Roberts et al., 2013; Yu et al., 2019b). Previously, however, it was not known how CEPs regulate seedling growth in foxtail millet (*Setaria italica*). In this study, we identified 14 *SiCEP* genes encoding 12 mature SiCEPs (Fig. S1B and Table S2). Expression patterns in tissues and in response to some abiotic stresses indicated that *SiCEP* genes play important roles in multiple development and abiotic stress responses (Fig. 2). Application of biosynthesized mature SiCEP3 and overexpression of *SiCEP3* revealed that *SiCEP3* negatively regulates plant growth in foxtail millet and *Arabidopsis* by promoting ABA transport and, to a lesser extent, ABA biosynthesis (Fig. 3A and B, Fig.4 A–C, and Fig. 5A and B). Finally, both SiCEP and ABA, alone or in combination, inhibit the kinase activities of CEPR1 and CEPR2. Collectively, these findings reveal crosstalk between CEP- and ABA-mediated abiotic stress responses in foxtail millet (*Setaria italica*), a novel C4 model plant (Yang et al., 2020).

Abiotic stress negatively impacts the growth and productivity of crop plants worldwide (Lamaoui et al., 2018). We found that *SiCEP* genes were significantly induced by ABA, PEG600, and salinity in both roots and shoots (Fig. 2), indicating the important roles of *SiCEP*s in response to salt and drought stresses in foxtail millet seedlings. These results were in agreement with previous reports implicating CEPs in abiotic stress responses in other species (Aggarwal et al., 2020; Stuhrwohldt and Schaller, 2019). Together, these data indicated that CEPs play conserved roles in regulating plant responses to abiotic stresses. Under abiotic stresses, ABA accumulates at high levels, which in turn inhibits primary root growth (Ma et al., 2019). We achieved effects similar to those of CEPs by applying SiCEP3 to foxtail millet seedlings or overexpressing the gene in hairy roots (Fig. 3A and B, Fig.4 A-C, and Fig.5 A and B). Root growth inhibition was aggravated when SiCEP3 concentration increased in the presence of ABA. Moreover, SiCEP3 activated the expression of genes encoding ABA transporters, such as *ABCG40* and *NRT1.2/NPF4.6,* in both foxtail millet and *Arabidopsis* seedlings (Fig. 3E and F, Fig. 4E and F, and Fig. 5D and E). These findings suggested that SiCEP3 promotes ABA import. Upregulation of *PYL4* and higher ABA content in both foxtail millet and *Arabidopsis* seedlings confirmed that ABA import was promoted by SiCEP3 (Fig. 3D and G, Fig.4 D and G, and Fig.5 C and F). Seedlings of the *nrt1.2abcg40* and *pyr1/pyl124* mutants had longer roots than wild-type seedlings (Fig. 5A and B), further mutant phenotype displayed and the requirement for *ABCG40*, *NRT1.2/NPF4.6,* and ABA receptors in SiCEP3-mediated promotion of ABA import and ABA signaling. Previously, we showed that NRT1.2 and PYL4 were modified post-translationally by CEPR2 (Yu et al., 2019b; Zhang et al., 2021). Therefore, CEPs regulate ABA import and receptors at both the transcriptional and post-translational levels.

Although most CEPs were mainly expressed in roots in rice (Sui et al., 2016), *Arabidopsis* (Ohyama et al., 2008), and foxtail millet seedlings (Fig. 2A), SiCEP9 was expressed at high levels in shoots (Fig.2A). Moreover, OsCEP6.1 is highly expressed in panicles in rice, resulting in inhibition of panicle and seed growth (Sui et al., 2016). Most *SiCEP* genes were transcribed at higher levels in panicles than in stems and leaves (Fig.2A). Three genes, including *SiCEP3*, *SiCEP4*, and *SiCEP5*, were expressed at higher levels in panicles than in roots (Fig.2A). These results suggested that SiCEPs play roles in regulating shoot growth, and panicle and seed development in foxtail millet plants. ABA plays a negative role in seed development (Cheng et al., 2014; Kang et al., 2013; Zegada-Lizarazu and Monti, 2019; Zhou et al., 2009). In *Arabidopsis*, ABA induces the expression of ABI5 gene, whose product inhibits the expression of *SHB1*, one of the key genes involved in positive regulation of seed development, leading to small seeds. Both AtCEPs (Delay et al., 2013) and SiCEP3 (Fig. 2C) could be transported to shoots, and possibly to panicle and developed seeds, indicating that both locally produced CEPs and peptides transported over long distances participate in the inhibition of shoot growth and panicle and seed development. Thus, CEPs play roles in the growth and development of multiple organs.

CEPs can balance plant growth and development and abiotic stress responses (Fig. 6B). Under normal conditions, low CEP levels are maintained by low expression of CEP genes (Fig. 2), resulting in activation of the CEPR1 and CEPR2 kinases (Fig. 6A). Active CEPR2 phosphorylates NRT1.2 and PYL4, promoting their degradation via UBC32-, UBC33-, and UBC34-mediated degradation (Yu et al., 2019b; Zhang et al., 2021). Consequently, ABA transport and signaling are inhibited, allowing plant growth. Under abiotic stresses, ABA and CEPs accumulate to high levels. ABA induces the expression of *ABCG40*, *NRT1.2* and *PYL4* genes (Fig. 3, 4, and 5), and both ABA and CEPs inhibit the kinase activities of CEPR1 and CEPR2 (Fig. 6A). Inactivation of CEPR2 results in activation of NRT1.2 and PYL4, which promote ABA transport and ABA binding, resulting in transduction of ABA signaling (Yu et al., 2019b; Zhang et al., 2021). However, to further elucidate CEP-mediated ABA transport, future studies should seek to identify the targets of CEPR1 and CEPR2, as well as novel ABA transporters.

## Author contributions

C.W. and C.Z. conceived the original experiments; L.Z., R.Y., Y.W. and Q.X. performed experiments, analyzed the data, made the Fig.s, and wrote the original article; G.Y., S.Z., J.H., and K.Y. provided suggestions; C.W. and C.Z. supervised and complemented the writing. All authors read and approved the final manuscript.

## Acknowledgments

This work was supported by the National Key R&D Program of China (2018YFD1000704, 2018YFD1000700), the Major Program of Shandong Province Natural Science Foundation (ZR2018ZB0212), and the Natural Science Foundation of China (Grant number 31970292). We thank Dr. Heribert Hirt (Center for Desert Agriculture, King Abdullah University of Science and Technology) for kindly providing pyr1/pyl1/pyl2/pyl4 mutant, and Prof. Yinggao liu (Shandong Agricultural University, China) for the *abi3*, *abi5*, *aba1*, *aba2* and *nced3* mutants.

## Data availability

All relevant data, vectors, and plant materials that support the findings of this study are available from the corresponding author upon request.

---

**Supplemental Figure 1. Gene and protein structures of SiCEPs in *Setaria italica*.**

(A) Gene structures of SiCEPs, created with GSDS.

(B) Signal peptide positions and CEP domains of SiCEPs. The scissors symbol indicates the cleavage site.

(C) Three-dimensional structures of SiCEPs predicted using the I-TASSER website.

**Supplemental Figure 2. Expression pattern analysis of SiCEPs.**

(A) Expression of SiCEPs in four different tissues.

(B) Cis-element analysis in the promoter regions 1500 bp upstream of SiCEPs.

(C) Statistic of cis-elements shown in (B).

**Supplemental Figure 3. Phenotypes of foxtail millet seedlings treated with SiCEP3 and ABA for 30 days.**

(A) Phenotypes of foxtail millet seedlings grown for 30 days on 1/2 MS (Control) or with 1 μM SiCEP3, 7.5 μM ABA, or the combination.

(B) Primary root length of foxtail millet seedlings shown in (A). Error bars indicate SEM (N = 3). Bars labeled with different lowercase letters are significantly different from one another (P < 0.05; one way ANOVA).

**Supplemental Figure 4. Responses of ABA response and ABA synthesis mutants to ABA or AtCEP1.**

(A) Phenotypes of WT, *abi3*, *abi5*, *aba1*, *aba2*, *nced3,* and *pyr1/pyl124* seedlings grown for 7 days on 1/2 MS containing 1 μM AtCEP1, 1 μM ABA, AtCEP1 +1 μM ABA, or 5 μM AtCEP1 +1 μM ABA.

(B) Lengths of primary roots of seedlings shown in (A). Error bars indicate SEM (N = 3). Bars labeled with different lowercase letters are significantly different from one another (P < 0.05; one way ANOVA).

**Table S1. Primers used in this study.**

**Table S2. Amino acid number, pI, molecular weight, signal peptide, and mature peptide of SiCEPs.**

